# Exploration of natural red-shifted rhodopsins using a machine learning-based Bayesian experimental design

**DOI:** 10.1101/2020.04.21.052548

**Authors:** Keiichi Inoue, Masayuki Karasuyama, Ryoko Nakamura, Masae Konno, Daichi Yamada, Kentaro Mannen, Takashi Nagata, Yu Inatsu, Kei Yura, Oded Béjà, Hideki Kandori, Ichiro Takeuchi

## Abstract

Microbial rhodopsins are photoreceptive membrane proteins utilized as molecular tools in optogenetics. In this paper, a machine learning (ML)-based model was constructed to approximate the relationship between amino acid sequences and absorption wavelengths using ~800 rhodopsins with known absorption wavelengths. This ML-based model was specifically designed for screening rhodopsins that are red-shifted from representative rhodopsins in the same subfamily. Among 5,558 candidate rhodopsins suggested by a protein BLAST search of several protein databases, 40 were selected by the ML-based model. The wavelengths of these 40 selected candidates were experimentally investigated, and 32 (80%) showed red-shift gains. In addition, four showed red-shift gains > 20 nm, and two were found to have desirable ion-transporting properties, indicating that they were potentially useful in optogenetics. These findings suggest that an ML-based model can reduce the cost for exploring new functional proteins.

## Introduction

Microbial rhodopsins are photoreceptive membrane proteins widely distributed in bacteria, archaea, unicellular eukaryotes, and giant viruses1. They consist of seven transmembrane (TM) α helices, with a retinal chromophore bound to a conserved lysine residue in the seventh helix (Fig. 1a). The first microbial rhodopsin, bacteriorhodopsin (BR), was discovered in the plasma membrane of the halophilic archaea *Halobacterium salinarum* (formerly called *H. halobium*)^2^. BR forms a purple-coloured patch in the plasma membrane called purple membrane, which outwardly transports H^+^ using sunlight energy^3^. After the discovery of BR, various types of microbial rhodopsins were reported from diverse microorganisms, and recent progress in genome sequencing techniques has uncovered several thousand microbial rhodopsin genes^1,4–6^. These microbial rhodopsins show various types of biological functions upon light absorption, leading to all-*trans*-to-13-*cis* retinal isomerization. Among these, ion transporters, including light-driven ion pumps and light-gated ion channels, are the most ubiquitous (Fig. 1b). Ion-transporting rhodopsins can transport several types of cations and anions, including H^+^, Na^+^, K^+^, halides (Cl^−^, Br^−^, I^−^), NO^−^, and SO4^2–1,7–9^. The molecular mechanisms of ion-transporting rhodopsins have been detailed in numerous biophysical, structural, and theoretical studies^1^.

**Fig. 1.**
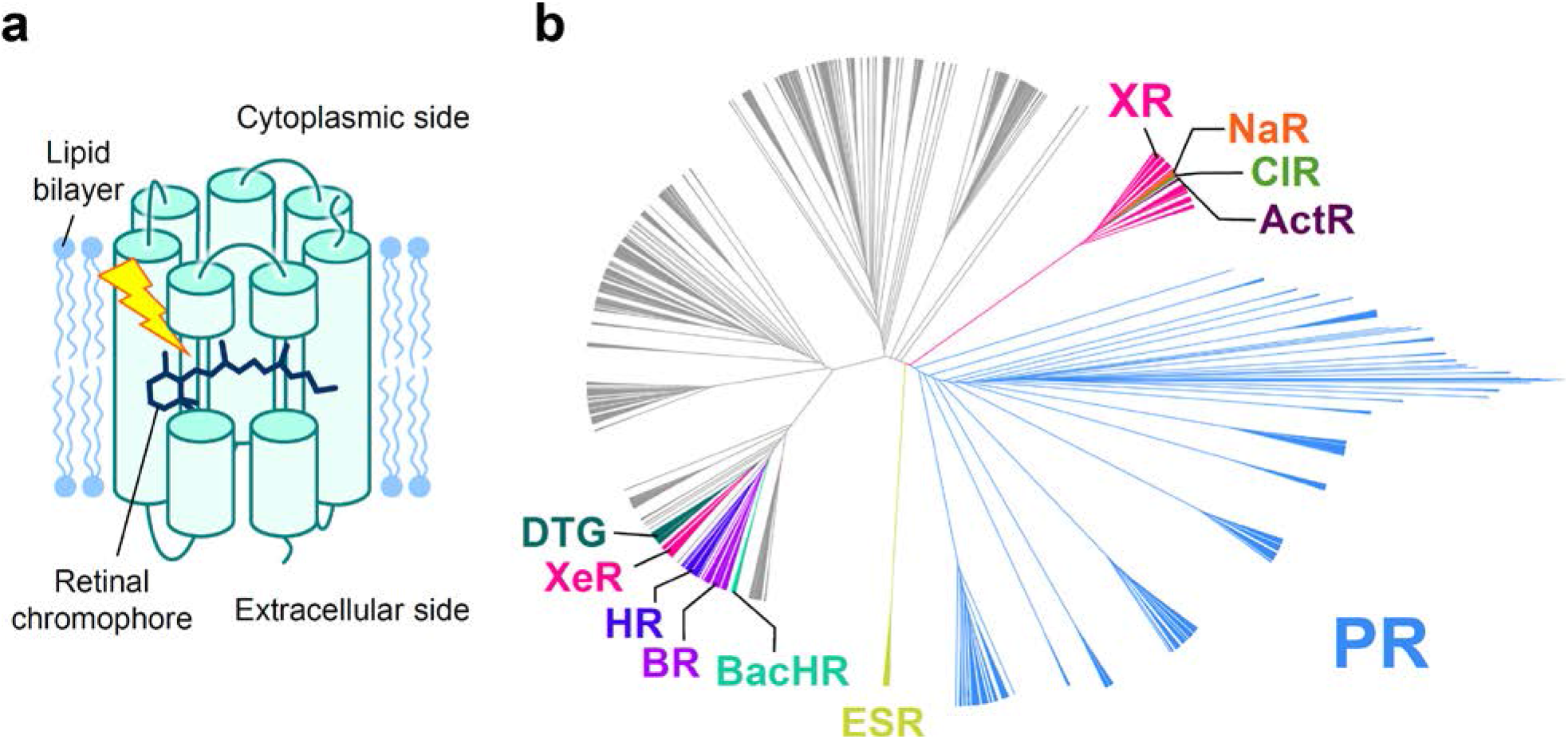
Structure and phylogenetic tree of microbial rhodopsins. **a** Schematic structure of microbial rhodopsins. b Phylogenic tree of microbial rhodopsins. The subfamilies of light-driven ion-pump rhodopsins targeted in this study are differently coloured; non-ion-pump microbial rhodopsins are shown in grey.

In recent years, many ion-transporting rhodopsins have been used as molecular tools in optogenetics to control the activity of animal neurons optically *in vivo* by heterologous expression^10^, and optogenetics has revealed various new insights regarding the neural network relevant to memory, movement, and emotional behaviour^11–14^. However, strong light scattering by biological tissues and the cellular toxicity of shorter wavelength light make precise optical control difficult. To circumvent this difficulty, new molecular optogenetics tools based on red-shifted rhodopsins that can be controlled by weak scattering and low toxicity longer-wavelength light are urgently needed. Therefore, many approaches to obtain red-shifted rhodopsins, including gene screening, amino acid mutation based on biophysical and structural insights, and the introduction of retinal analogs, have been reported^15–17^ Recently, a new method using a chimeric rhodopsin vector and functional assay was reported to screen the absorption maximum wavelengths (*Λ*_max_) and proton transport activities of several microbial rhodopsins present in specific environments18. This method identified partial sequences of red-shifted yellow (560–570 nm)-absorbing proteorhodopsin (PR), the most abundant outward H^+^-pumping bacterial rhodopsin subfamily, from the marine environment. Although these works identified several red-shifted rhodopsins^14,15,17,19^, those showing ideally red-shifted absorption and high ion-transport activity sufficient for optical control *in vivo* have yet to be obtained.

As an alternative approach, we recently introduced a data-driven machine learning (ML)- based approach^20^. In the previous study, we demonstrated how accurately the absorption wavelength of rhodopsins could be predicted based on the amino acid types on each position of the seven TM helices^20^. We constructed a database containing 796 wild-type (WT) rhodopsins and their variants, the *λ*_max_ of which had been reported in earlier studies. Then, we demonstrated the prediction performance of the ML-based prediction model using a data-splitting approach, i.e., the data set was randomly divided into a training set and a test set; the former was used to construct the prediction model, and the latter was used to estimate the prediction ability. The results of this “proof-of-concept” study suggested that the absorption wavelengths of an unknown family of rhodopsins could be predicted with an average error of ±7.8 nm, which is comparable to the mean absolute error of *λ*_max_ estimated by the hybrid quantum mechanics/molecular mechanics (QM/MM)^21^ method. Considering the computational cost of both approaches, the ML-based approach is much more efficient than QM/MM approach, while the latter provides insights on the physical origin controlling *λ*_max_.

Encouraged by this result, in this study, we used an ML-based approach to screen more red-shifted rhodopsins from among 3,064 new candidates collected from public databases (non-redundant and metagenomic rhodopsin genes from the National Center for Biotechnology Information [NCBI] and *Tara* Oceans data sets) for which the absorption wavelengths have not been investigated. The goal of the present study was to identify rhodopsins with a *λ*_max_ longer than the wavelengths of representative rhodopsins in each subfamily of microbial rhodopsins for which the *λ*_max_ has already been reported (base wavelengths). Here, we call the red-shift change in the wavelength from the base wavelength the “red-shift gain”. We focus on the problem of identifying rhodopsins with large red-shift gains because this would lead to the identification of amino acid types and residue positions that play important roles in red-shifting absorption wavelengths. In addition, in optogenetics applications, it is practically important to have a wide variety of ion-pumping rhodopsins from each subfamily to construct a new basis for rhodopsin toolboxes with red-shifted absorption and various types of ion species that can be transported. To screen rhodopsins that would have large red-shift gains, it is necessary to consider the uncertainty of prediction in the form of “predictive distributions”^22^. By using predictive distributions, it is possible to consider appropriately the “exploration–exploitation trade-off” in screening processes^23,24^, where exploration indicates an approach that prefers candidates with larger predictive variances, and exploitation indicates an approach that prefers candidates with longer predictive mean wavelengths (Fig. 2). In this paper, we employ a Bayesian modeling framework to compute the predictive distributions of candidate rhodopsin red-shift gains. We then consider an exploration–exploitation trade-off by selecting candidate rhodopsins based on a criterion called “expected red-shift gains”.

**Fig. 2.**
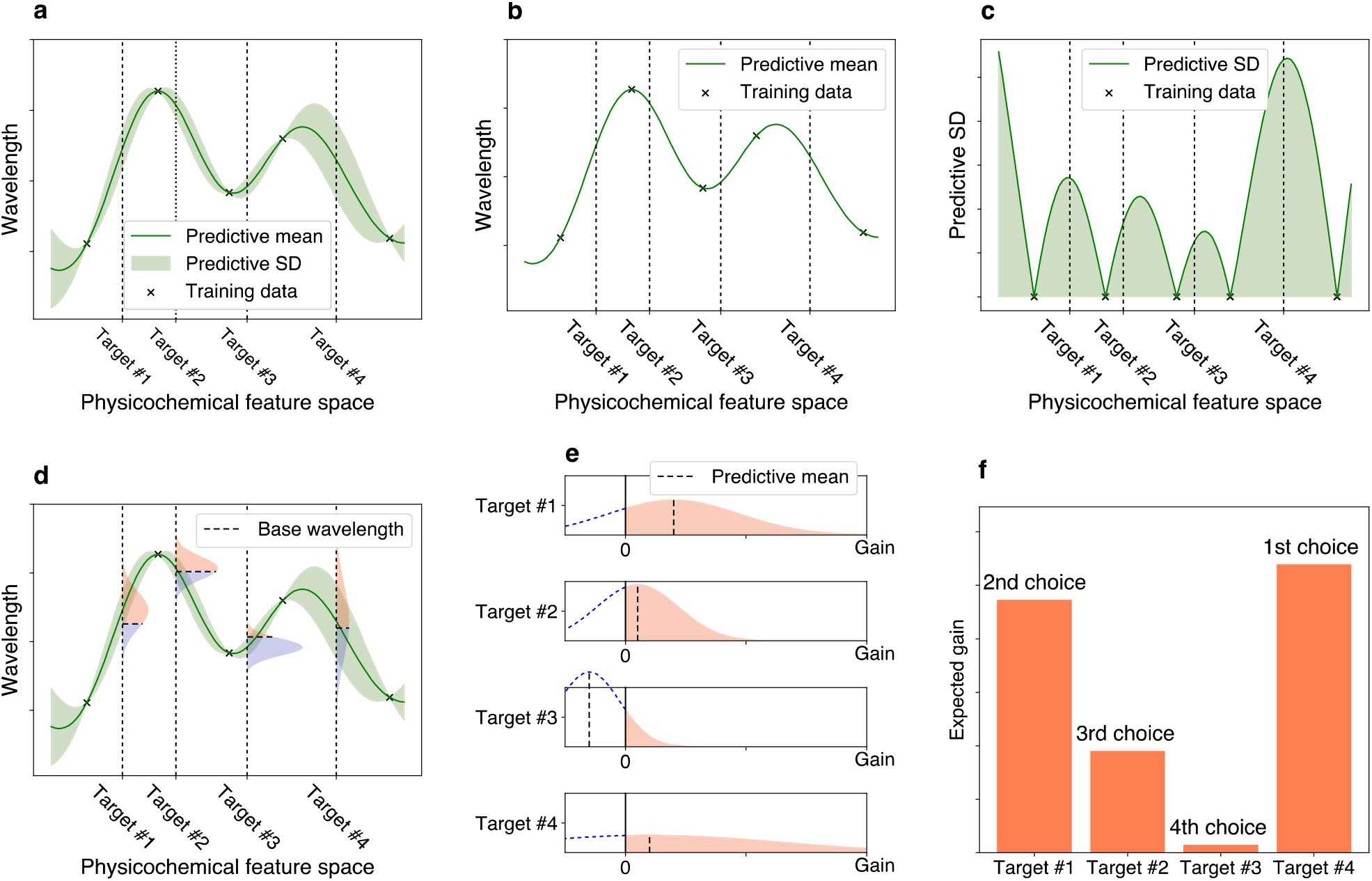
Illustrations of exploration–exploitation for screening rhodopsins with red-shift gain. **a** Bayesian prediction model constructed using the current training data (black crosses). The prediction model is represented by the predictive mean and predictive standard deviation (SD). The horizontal axis schematically illustrates the space of proteins defined through physicochemical features. The four vertical dotted lines indicate target proteins (candidates to synthesize). **b** Predictive mean. This function is defined as the expected value of the probabilistic prediction by the Bayesian model. **c** Predictive SD. Since the predictive SD represents the uncertainty of the prediction, it has a larger value when the training data points do not exist nearby. **d** The distributions on the vertical dotted lines represent the predictive distributions, and the horizontal dashed lines are the base wavelengths of the target points. The base wavelength is different for each target point because it depends on the subfamily of the protein. **e** The density of the predictive distribution of each target protein on its red-shift gain value. The gain is defined as the predicted wavelength subtracted by the base wavelength, and if it is negative, the value is truncated as 0. This can be seen as a “benefit” that can be obtained by observing the target protein. *f* Expected value of the red-shift gain. This provides a ranking list from which the next candidates to be experimentally investigated can be determined. Target #4 has the largest expected gain, although target #1 has the largest increase in the predictive mean compared with base wavelength in e. Because of its larger SD (as shown in a, c, d, and e), target #4 is probabilistically expected to have a larger gain than the other targets.

In this paper, we updated the ML-based model used in our previous study^20^ so that it could properly compute expected red-shift gains and applied this new model to 3,064 ion-pumping rhodopsin candidates derived from archaeal and bacterial origins that can be easily expressed in *Escherichia coli* (Fig. 1b). We then selected 66 candidates for which the expected gains were > 10 nm, and experimentally investigated their wavelengths by introducing the synthesized rhodopsin genes into *E. coli*. Of these 66 selected candidates, 40 showed significant colouring in *E. coli* cells, 32 showed actual red-shift gains, seven showed blue-shifts, and one showed no change, suggesting that our ML-based model enables more efficient screening of red-shifted rhodopsin genes compared with random choice (i.e., 80.5% [32/40] of the selected candidates showed red-shift gains with *p* < 10^−3^ in a binomial test). We then investigated the ion-transportation properties of the rhodopsins whose red-shift gains were > 20 nm, and found that some actually had desired ion-transporting properties, suggesting that they (and their variants) could potentially be used as new optogenetics tools. Furthermore, the differences in the amino-acid sequences of the newly examined rhodopsins and the representative ones in the same subfamily could be used for further investigation of the red-shifting mechanisms.

## Results

### Construction of an ML-based model for computing expected red-shift gain

To compute the expected red-shift gains of a wide variety of rhodopsins, we updated various aspects of the ML model used in our previous study^19^. Figure 3 shows a schematic of the updating procedure. First, we added 97 WT microbial rhodopsins and their variants for which the *λ*_max_ had recently been reported in the literature or determined by our experiments, to a previously reported data set^20^. In other words, the new training data set consisted of the amino acid sequences and *λ*_max_ of 893 WT microbial rhodopsins and their variants (Extended Data Table 1). Second, the new ML model used only *N* = 24 residues located around the retinal chromophore (Extended Data Figure 1) because our previous study^19^ indicated that amino acid residues at these 24 positions play significant roles in predicting absorption wavelengths (Fig. 3a). Third, *M =* 1818 amino acid physicochemical features (Extended Data Table 2) were used as inputs in the ML model, as opposed to the amino acid types used in the previous ML model. This enabled us to predict the absorption wavelengths of a wide range of target rhodopsins that contain unexplored amino acid types in the training data at certain positions.

**Fig. 3.**
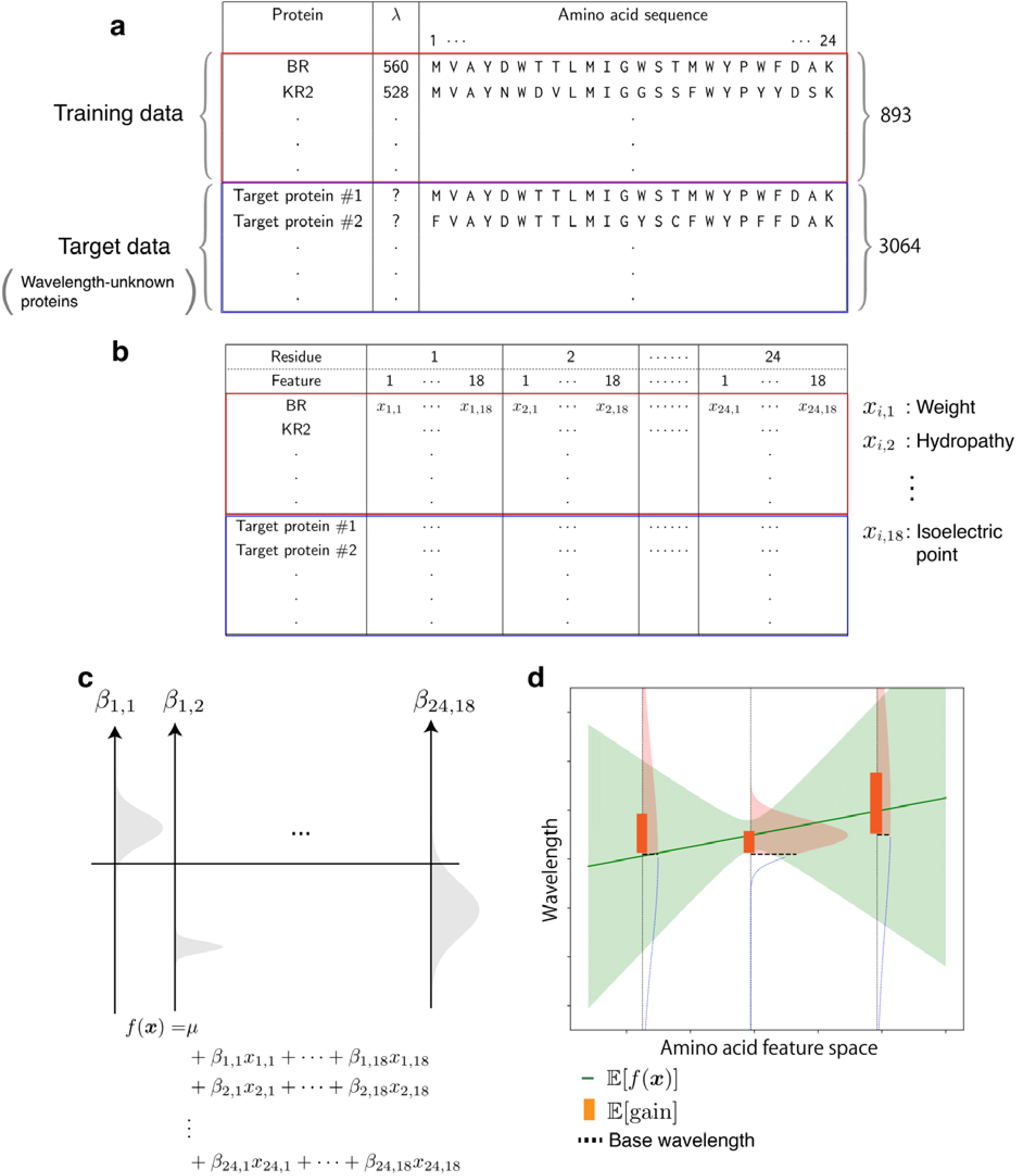
Overview of the ML-based exploration of natural red-shifted rhodopsins. **a** Using existing experimental data, a training data set consisting of pairs of a wavelength *λ*_max_ and an amino acid sequence was constructed. A particular focus was placed on the 24 amino acid residues around the retinal chromophore to build an ML-based prediction model. A set of protein sequences with no known wavelength was also collected as target proteins. **b** All amino acid sequences were transformed into physicochemical features, leading to 24 × 18 = 432 dimensional numerical representations of each protein. **c** A linear regression model was constructed using the Bayesian approach. Each regression coefficient *β_i,j_* was estimated as a distribution (shown as a gray region). The broadness of these distributions represent the uncertainty of the current estimation. **d** The expected red-shift gain values were evaluated for the target proteins. The green region is the standard deviation of the prediction. The red shaded region in the vertical distribution corresponds to the probability that the wavelength is larger than the base wavelength (dashed line), which is determined by the subfamily of the microbial rhodopsin. The bar represents the expected red-shift gain, defined by the expected value of the increase from the base wavelength.

Therefore, an amino acid sequence is transformed into an *M* × *N* = 432 dimensional feature vector ***x*** ∈ ℝ^*MN*^ by concatenating *x_i,j_*, the *j*-th feature of the *i*-th residue (Fig. 3b). We consider a linear prediction model 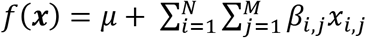, where *β_i,j_* is the parameter for the *j*-th feature of the *i*-th residue, and *μ* is the intercept term.

Finally, to consider the exploration–exploitation trade-off appropriately in the screening process, we introduce a Bayesian modeling framework, which allows us to compute the predictive distributions of red-shift gains. Specifically, we employed Bayesian sparse modeling called BLASSO^25^ (see the Methods section for details). This enables us to provide not only the mean, but also the variance of the predicted wavelengths. Unlike classical regression analysis, BLASSO regards the model parameters *β_i,j_* and *μ* as random variables generated from underlying distributions, as illustrated in Figure 3c. Therefore, the wavelength prediction *f*(***x***) is also represented as a distribution. The red-shift gain is defined as gain = max(*f*(***x***) − *λ*_base_, 0), where *λ*_base_ is the wavelength of the representative rhodopsin in the same subfamily whose *λ*_max_ has been experimentally determined and reported in the literature (Extended Data Table 3). Note that the red-shift gain is positive if *f*(***x***) is greater than *λ*_base_; otherwise, it takes the value of zero. Since *f*(***x***) is regarded as a random variable in BLASSO, the red-shift gain is also regarded as a random variable. Therefore, we employ the expected value of the redshift gain, denoted by 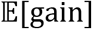, as the screening criterion where 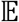 represents the expectation of a random variable. Illustrative examples of 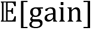 are shown in Figure 3d. Unlike the simple expectation of the wavelength prediction 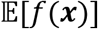, 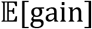 depends on the variance of the predictive distribution. This encourages the exploration of rhodopsin candidates having large uncertainty (for exploration), as opposed to only those having longer wavelengths with high confidence (for exploitation).

### Screening potential red-shifted microbial rhodopsins based on expected red-shift gains

The target data set to explore red-shifted microbial rhodopsins was constructed by collecting putative microbial rhodopsin genes collected by a protein BLAST (blastp) search^26^ of the NCBI non-redundant protein and metagenome databases^27^, as well as the *Tara* Oceans microbiome and virome databases^28^. As a result, we obtained a non-redundant data set of 5,558 microbial rhodopsin genes (Fig. 1b). The sequences were aligned by ClustalW and categorized to subfamilies of microbial rhodopsins based on the phylogenic distances, as reported previously^29^. Among these, 3,064 rhodopsin genes from bacterial and archaeal origins were extracted because their *λ*_max_ can be easily measured by expressing in *E. coli* cells. We calculated the 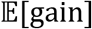[gain] of these 3,064 genes (Extended Data Table 4), and then selected 66 genes of putative light-driven ion pump rhodopsins showing an 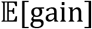 > 10 nm for further experimental evaluation, as ion pump rhodopsins can be used as new optogenetics tools.

### Experimental measurement of the absorption wavelengths of microbial rhodopsins showing high red-shift gains

We synthesized the selected 66 genes that showed an 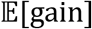 > 10 nm. These were then introduced into *E. coli* cells, and the proteins expressed in the presence of 10 μM *all-trans* retinal. As a result, 40 *E. coli* cells showed significant colouring, indicating significant expression of folded protein, and their *λ*≡ were determined by observing ultraviolet (UV)-visible absorption changes upon bleaching of the expressed rhodopsins through a hydrolysis reaction of their retinal with hydroxylamine, as previously reported^20^ (Fig. 4). The observed gains were compared with the 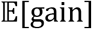 shown in Table 1. A full list of unexpressed genes is shown in Extended Data Table 5. In total, 32 of 40 genes showed a longer wavelength than their base wavelength (that is, positive red-shift gain) (Fig. 5), suggesting that our ML-based model can significantly improve the efficiency of screening to explore new red-shifted microbial rhodopsins compared with random sampling (*p* < 0.0002).

**Fig. 4.**
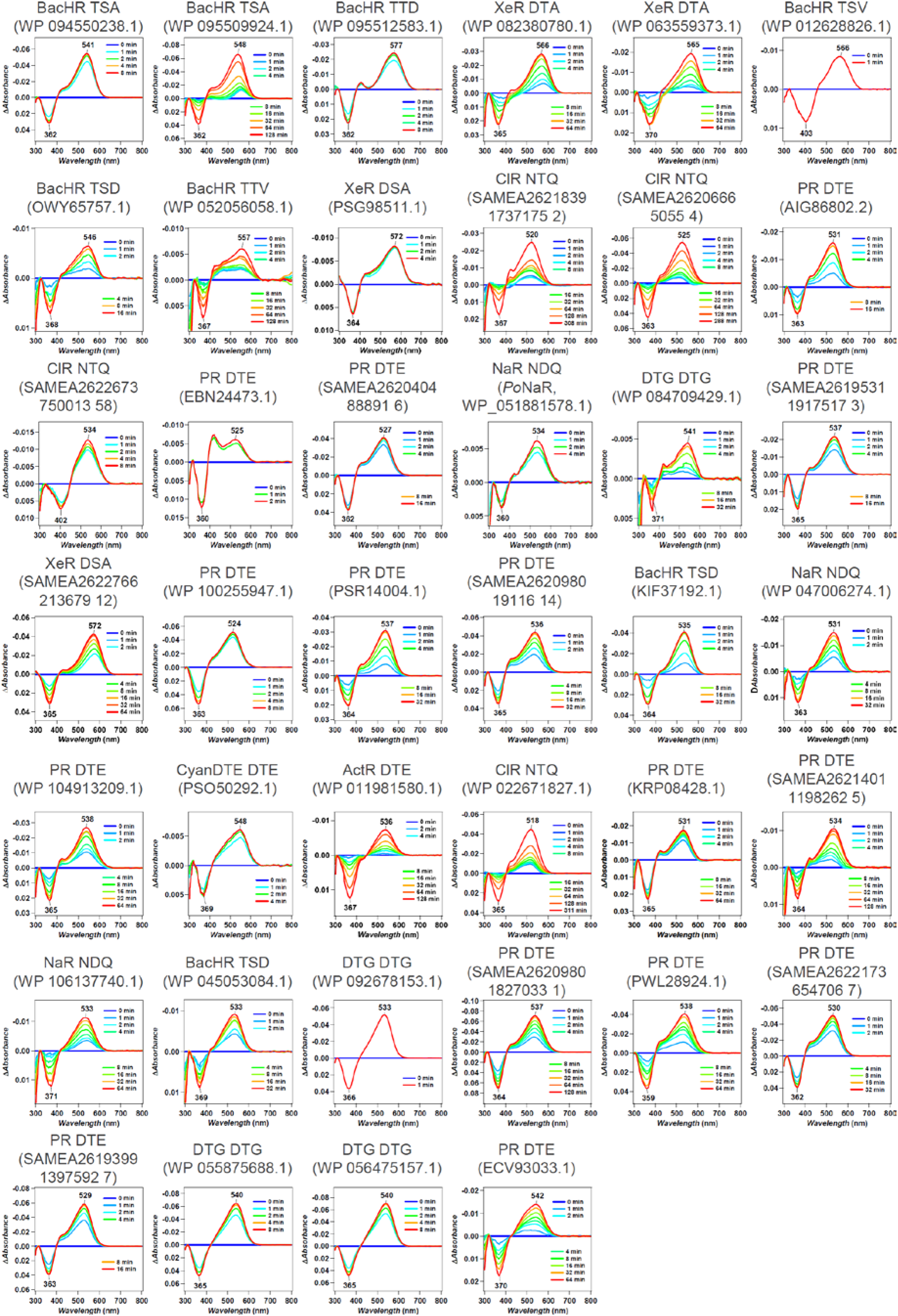
*λ*_max_ of 40 microbial rhodopsins in solubilized *E. coli* membrane observed upon hydroxylamine bleach reaction. The difference absorption spectra between before and after hydroxylamine bleaching reaction of microbial rhodopsins in solubilized *E. coli* membrane. The *λ*_max_ of each rhodopsin was determined by the peak positions of the absorption spectra of the original proteins, and the absorption of retinal oxime produced by the reaction of retinal Schiff base and hydroxylamine was observed as a negative peak at around 360–370 nm.

**Table 1.** Predicted and observed gains of 40 microbial rhodopsins expressed in *E. coli*.Figures

| Origin | Accession | Subfamily | Motif | Base wavelength<br>/ nm | E [gain] | Observed<br>wavelength<br>/ nm | (Observed<br>wavelength) –<br>(base wavelength) /<br>nm |
| --- | --- | --- | --- | --- | --- | --- | --- |
| *Rubricoccus marinus* | WP 094550238.1 | BacHR | TSA | 537 | 40.7 | 541 | 4 |
| *Rubrivirga marina* | WP 095509924.1 | BacHR | TSA | 537 | 39.8 | 548 | 11 |
| *Rubrivirga marina* | WP 095512583.1 | BacHR | TTD | 537 | 35.5 | 577 | 40 |
| *Bacillus* sp. CHD6a | WP 082380780.1 | XeR | DTA | 565 | 35.3 | 566 | 1 |
| *Bacillus horikoshii* | WP 063559373.1 | XeR | DTA | 565 | 35.3 | 565 | 0 |
| *Cyanothece* sp. PCC 7425 | WP 012628826.1 | BacHR | TSV | 537 | 32.9 | 566 | 29 |
| *Cyanobacterium* TDX16 | OWY65757.1 | BacHR | TSD | 537 | 32.9 | 546 | 9 |
| *Myxosarcina* sp. GI1 | WP 052056058.1 | BacHR | TTV | 537 | 31.2 | 557 | 20 |
| *Nanohaloarchaea*<br>archaeon SW 7 43 1 | PSG98511.1 | XeR | DSA | 565 | 29.2 | 572 | 7 |
| Metagenome sequence | SAMEA2621839 1737175<br>2 | CIR | NTQ | 530 | 25.7 | 520 | -10 |
| Metagenome sequence | SAMEA2620666 5055 4 | CIR | NTQ | 530 | 25.1 | 525 | -5 |
| *Nonlabens* sp. YIK11 | AIG86802.2 | PR | DTE | 520 | 21.5 | 531 | 11 |
| Metagenome sequence | SAMEA2622673 750013<br>58 | CIR | NTQ | 530 | 21.4 | 534 | 4 |
| Metagenome sequence | EBN24473.1 | PR | DTE | 520 | 20.0 | 525 | 5 |
| Metagenome sequence | SAMEA2620404 88891 6 | PR | DTE | 520 | 20.0 | 527 | 7 |
| *Parvularcula oceani* | WP_051881578.1 | NaR | NDQ | 525 | 19.7 | 534 | 9 |
| *Rubrobacter aplysinae* | WP 084709429.1 | DTG | DTG | 535 | 19.5 | 541 | 6 |
| Metagenome sequence | SAMEA2619531 1917517<br>3 | PR | DTE | 520 | 18.0 | 537 | 17 |
| Metagenome sequence | SAMEA2622766 213679<br>12 | XeR | DSA | 565 | 17.8 | 572 | 7 |
| *Reinekea forsetii* | WP 100255947.1 | PR | DTE | 520 | 17.1 | 524 | 4 |
| *Bacteroidetes* bacterium | PSR14004.1 | PR | DTE | 520 | 15.4 | 537 | 17 |
| Metagenome sequence | SAMEA2620980 19116<br>14 | PR | DTE | 520 | 15.4 | 536 | 16 |
| *Hassallia byssoidea*<br>VB512170 | KIF37192.1 | BacHR | TSD | 537 | 15.1 | 535 | -2 |
| *Erythrobacter*<br>*gangjinensis* | WP 047006274.1 | NaR | NDQ | 525 | 13.7 | 531 | 6 |
| *Pontimonas salivibrio* | WP 104913209.1 | PR | DTE | 520 | 12.2 | 538 | 18 |
| *Cyanobacteria* bacterium<br>QH 1 48 107 | PSO50292.1 | CyanDTE | DTD | 545 | 12.0 | 548 | 3 |
| *Kineococcus radiotolerans* | WP 011981580.1 | ActR | DTE | 540 | 11.2 | 536 | -4 |
| *Sphingopyxis*<br>*baekryungensis* | WP 022671827.1 | CIR | NTQ | 530 | 11.0 | 518 | -12 |
| *Sphingobacteriales*<br>bacterium BACL12<br>MAG120802bin5 | KRP08428.1 | PR | DTE | 520 | 10.9 | 531 | 11 |
| Metagenome sequence | SAMEA2621401 1198262<br>5 | PR | DTE | 520 | 10.9 | 534 | 14 |
| *Spirosoma oryzae* | WP 106137740.1 | NaR | NDQ | 525 | 10.8 | 533 | 8 |
| *Aliterella atlantica* | WP 045053084.1 | BacHR | TSD | 537 | 10.8 | 533 | -4 |
| *Rosenbergiella nectarea* | WP 092678153.1 | DTG | DTG | 535 | 10.8 | 533 | -2 |
| Metagenome sequence | SAMEA2620980 1827033<br>1 | PR | DTE | 520 | 10.4 | 537 | 17 |
| *Fluviicola* sp. XM24bin1 | PWL28924.1 | PR | DTE | 520 | 10.4 | 538 | 18 |
| Metagenome sequence | SAMEA2622173 654706<br>7 | PR | DTE | 520 | 10.4 | 530 | 10 |
| Metagenome sequence | SAMEA2619399 1397592<br>7 | PR | DTE | 520 | 10.4 | 529 | 9 |
| *Sphingomonas* sp. Leaf34 | WP 055875688.1 | DTG | DTG | 535 | 10.3 | 540 | 5 |
| *Sphingomonas* sp. Leaf38 | WP 056475157.1 | DTG | DTG | 535 | 10.3 | 540 | 5 |
| Metagenome sequence | ECV93033.1 | PR | DTE | 520 | 10.3 | 542 | 22 |

**Fig. 5.**
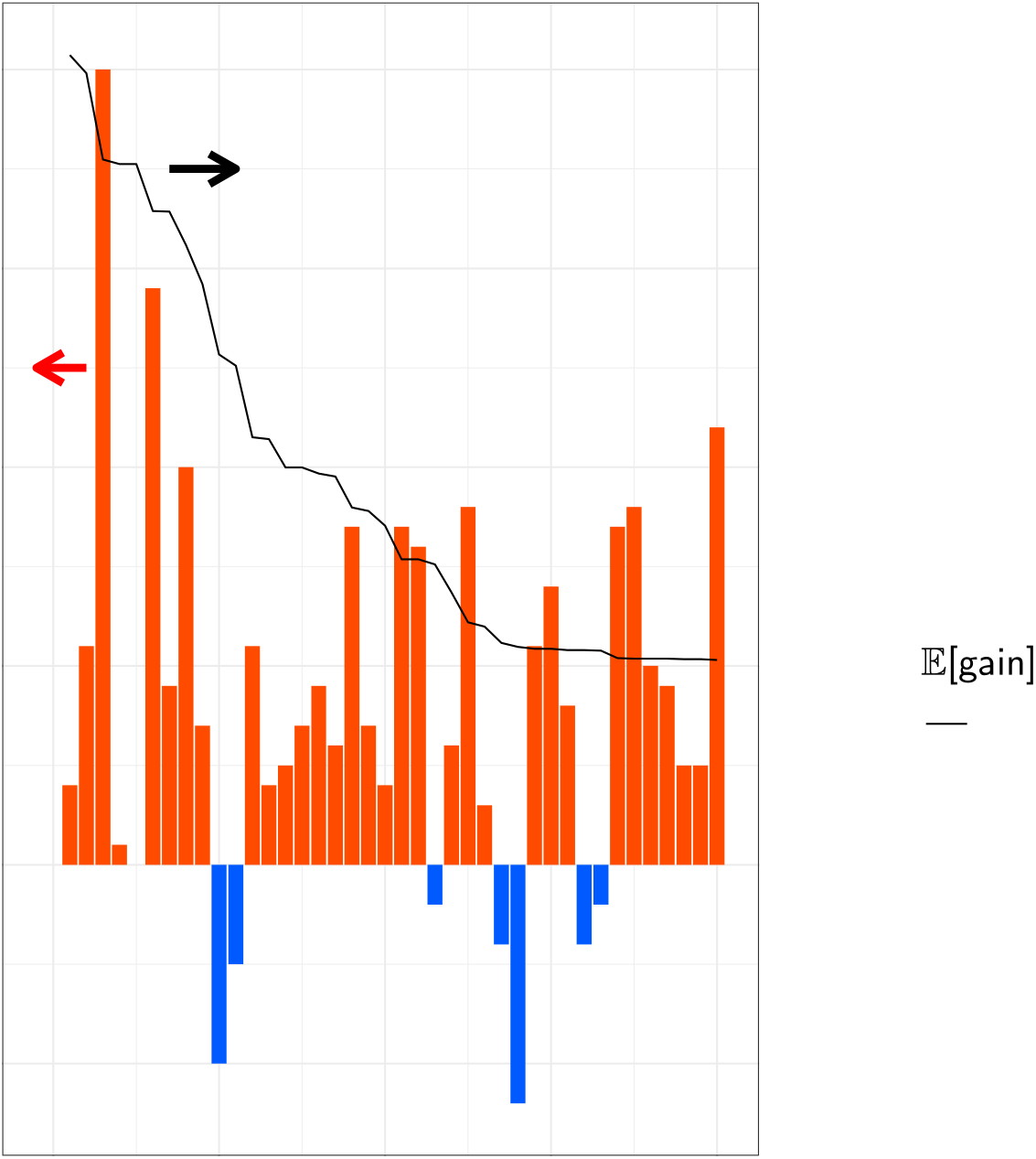
Observed wavelengths and expected red-shift gains. The predicted and observed red-shift (and blue-shift) gains for the 40 candidate rhodopsins that showed significant coloring in *E. coli* cells. Differences between observed and base wavelengths are shown by the bars. The red bars indicate red-shift from the base wavelength, while the blue bars indicate observed wavelengths that were shorter than the base wavelengths. Proteins are sorted in the descending order by 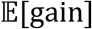, as shown by the black line. Among the 40 candidates, 33 (82.5%) showed red-shift gains, suggesting that the proposed ML-based model can screen red-shifted rhodopsins more efficiently than random choice.

### Ion-transport function of red-shifted microbial rhodopsins

Overall, four of the 40 rhodopsins showed red-shifted absorption > 20 nm compared with the base wavelengths (Table 1): three were halorhodopsins (HRs) from bacterial species^9,30,31^ (to distinguish classical HRs from archaeal species, these are hereafter referred to as bacterial-halorhodopsins [BacHRs]), and one was a PR^32^. Their ion-transport activities were then investigated by expressing in *E. coli* cells and observing the pH change in external solvent (Fig. 6). Upon light illumination, BacHRs from *Rubrivirga marina* and*Myxosarcina* sp. GI1 showed significant alkalization of external solvent, which was enhanced by addition of the protonophore (CCCP), which increases the H^+^ permeability of the cell membrane, and the light-dependent alkalizations disappeared when anions were exchanged from Cl^−^ to SO_4_^2−^, indicating that these were light-driven Cl^−^ pumps, similar to other rhodopsins in the same BacHR subfamily^9,30^. By contrast, *Cyanothece* sp. PCC 7425 did not show any significant transport. While no transporting function can be attributed to the heterologous expression in *E. coli*, it would have considerably different molecular properties from other BacHRs. PRs from a metagenome sequence (ECV93033.1) showed acidification of external solvent that was abolished by the addition of CCCP and was independent from ionic species in the solvent. Hence, this was a new red-shifted outward H^+^ pump compared with typical PRs whose *λ*_max_ are present at ca. 520 nm32. These light-driven ion-pumping rhodopsins with red-shifted *λ*_max_ have the potential to be applied as new optogenetics tools, and thus, warrant further study in the near future.

**Fig. 6.**
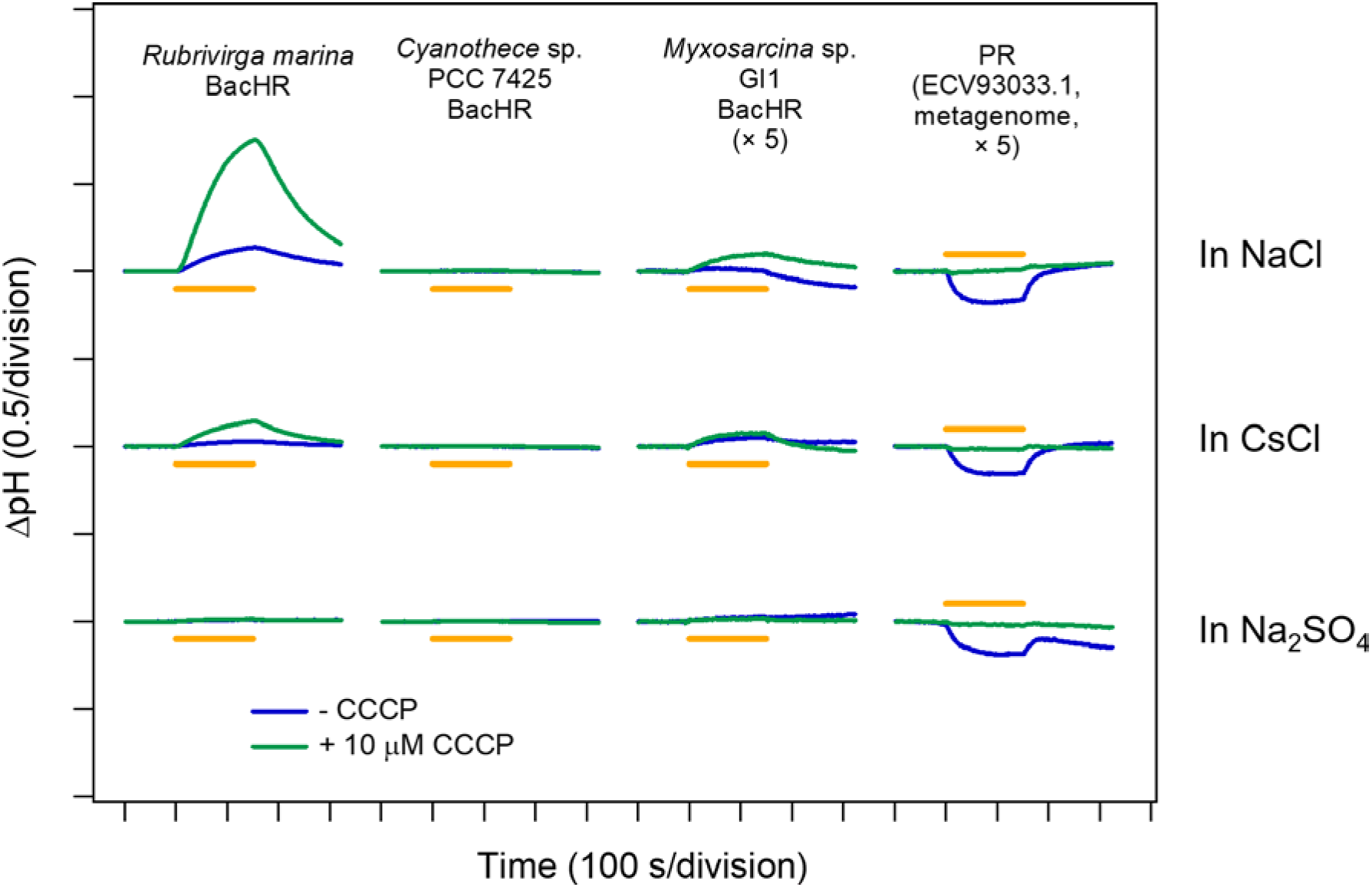
Light-driven ion-transport activities of microbial rhodopsins showed longer *λ*_max_. The light-induced pH change in the external solvent of *E. coli* cells expressing four microbial rhodopsins that showed a *λ*_max_ >20 nm longer than the base wavelength of the subfamily. The data obtained without and with 10 μM CCCP are indicated by the blue and green lines, respectively, in 100 mM NaCl (top), CsCl (middle), and Na2SO4 (bottom). Light was illuminated for 150 s (yellow solid lines).

## Discussion

Microbial rhodopsins show a wide variety of *λ*_max_ by changing steric and electrostatic interactions between all-*trans* retinal chromophores and surrounding amino acid residues. An understanding of the colo<u>u</u>r-tuning rule enables more efficient screening and the design of new red-shifted rhodopsins that have value as optogenetics tools, and our ML-based data-driven approach therefore provides a new basis to identify colour-regulating factors without assumptions.

We previously demonstrated that an ML-based model based on ~800 experimental results could predict the *λ*_max_ of microbial rhodopsins with an average error of ±7.8 nm. Encouraged by this result, in the present study, we constructed a new ML-based model to compute expected red-shift gains for a wide range of unknown families of microbial rhodopsins. As a result, 33 of 40 microbial rhodopsins were found to have red-shifted absorption compared with the base wavelengths of each subfamily of microbial rhodopsins (Table 1), suggesting that our data-driven ML approach can screen red-shifted microbial rhodopsin genes more efficiently than random choice.

By considering the exploration–exploitation trade-off, that is, to consider not only the expected value of the prediction, but also the uncertainty, it was possible to construct a red-shift protein screening process, as shown in Figure 7. Figure 7a shows the relationships between the prediction uncertainty (as measured by the standard deviation) and the observed red-shift gains. It can be seen that rhodopsins with red-shift gain are found in areas of not only low (small standard deviation), but also high prediction uncertainty (large standard deviation). Figure 7b shows the two-dimensional projection of the *d* = 432 dimensional feature space by principal component analysis. It can be seen that red-shift gains (red) are found for target proteins not only close to training proteins (green), but also far from training proteins. Figure 8 shows that the observed wavelengths and red-shift gains tend to be smaller than the predicted ones. We conjecture that these differences between the observed and predicted wavelengths and red-shift gains are due to modeling errors, possibly caused by a lack of sufficient information (e.g., three-dimensional structures) and modeling flexibility (e.g., nonlinear effects); in other words, rhodopsins having high prediction values partly by modeling errors have a high chance of being selected. Therefore, it would be valuable to develop a statistical methodology to eliminate selection bias due to modeling errors.

**Fig. 7.**
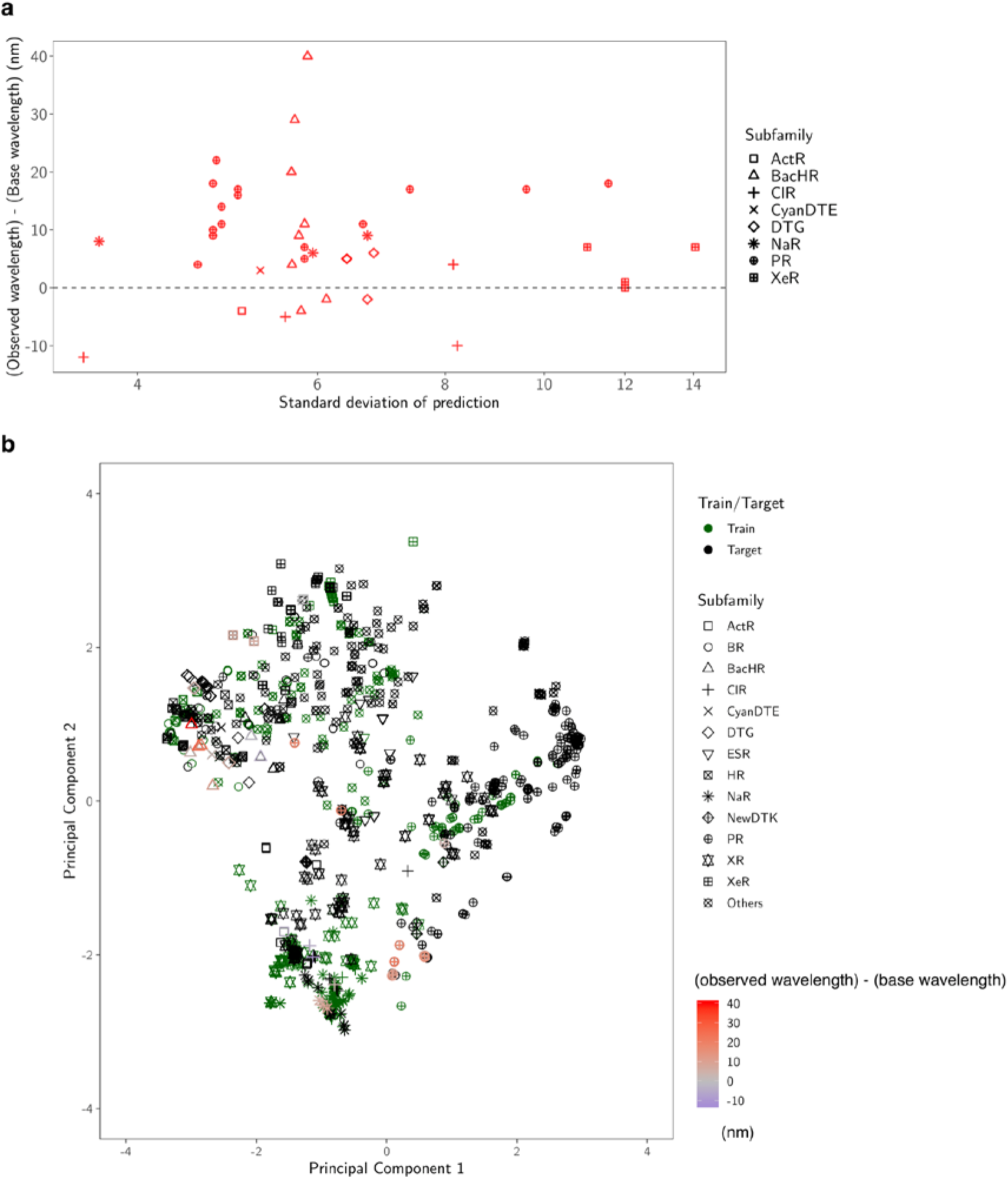
Diversity of the selected proteins. **a** Predicted standard deviation (horizontal axis) vs. observed gain (vertical axis). The marker shape represents the subfamily of each protein. **b** Two-dimensional projection created by principal component analysis. The original *d* = 432 dimensional feature space is projected onto the first two principal component directions. The first component (horizontal axis) explains 33% of the total variance of the original space, and the second (vertical axis) explains 17%. The green markers are the training data, and the black markers are the target data. For the synthesized proteins, differences in the observed and base wavelengths are shown by the color map. The results indicate that, by considering the exploration–exploitation trade-off, it was possible to make a red-shift protein screening process that considered not only the expected value of the prediction, but also the uncertainty.

**Fig. 8.**
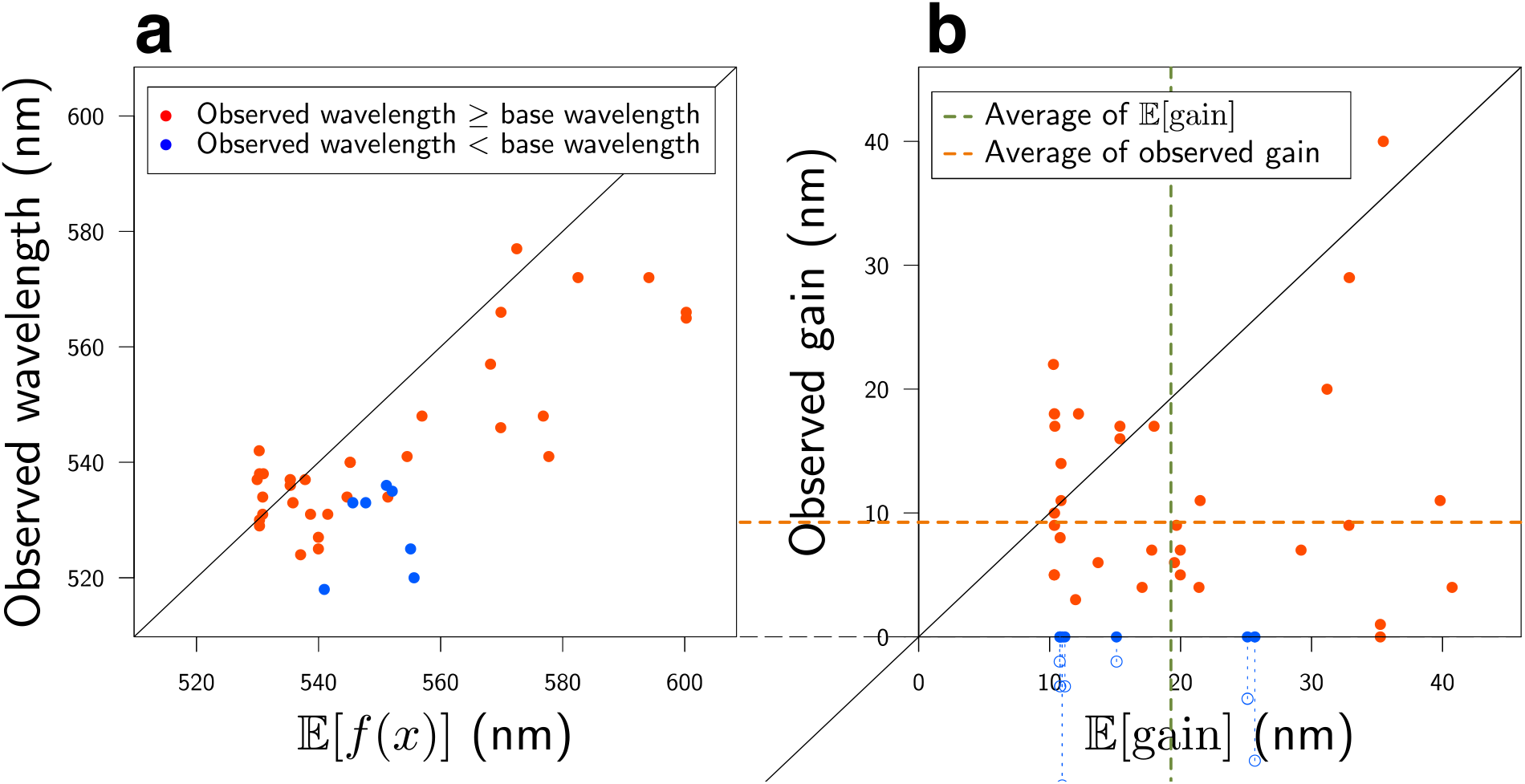
Comparisons of experimental observations and ML predictions. In these two plots, the red points have longer observed wavelengths than the base wavelength λ_base_, while the blue points have shorter observed wavelengths than *λ*_base_. a ML-based prediction of *λ*_max_ (horizontal axis) vs. experimentally observed *λ*_max_ (vertical axis). **b** Expected red-shift gain (horizontal axis) vs. observed gain (vertical axis). Since we selected rhodopsins having expected red-shift gains of > 10 nm, all the points on the horizontal axis are > 10 nm. The observed gain, defined by max(*λ*_max_ − *λ*_base_, 0), is nonnegative by definition. The green and orange dashed lines are the averages of the horizontal and vertical axes (19.3 nm and 9.3 nm), respectively. The results indicate that the observed wavelengths and red-shift gains tended to be smaller than the predicted ones. We conjecture that these differences between the observed and predicted wavelengths are due to modelling errors (see the Discussion for details).

Four rhodopsins showed red-shifted absorption > 20 nm than the base wavelength, three of which showed light-driven ion-transport function. Interestingly, while one BacHR from *Rubrivirga marina* (accession No.: WP 095512583.1) showed a 40-nm longer *λ*_max_ (577 nm) than the base wavelength, another 11-nm red-shifted BacHR (WP 095509924.1) was also identified from the same bacteria (Table 1). These BacHRs are highly similar to each other (55.2% identity and 70.6% similarity), and only four of 24 amino acid residues around the retinal chromophore differ. Hence, *R. marina* evolved two BacHRs with 29-nm different *λ*_max_ by small amino acid replacement; the amino acid residue(s) responsible for this color-tuning should be investigated in the future.

The differences in amino acids in three of 24 retinal-surrounding residues are known to play a color-tuning role in natural rhodopsins without affecting their biological function. These correspond to positions 93, 186, and 215 in BR (BR Leu93, Pro186, and Ala215, respectively)^16^. Position 93 is known to be diversified in the PR family (the well-known position 105 in PRs). Green-light-absorbing PRs (GPRs) have leucine as a BR, whereas glutamine is conserved in blue-light-absorbing PRs^4,18^. This colour-tuning effect by the difference between leucine and glutamine is known as the “L/Q-switch”^33^. Interestingly, while 29.8% of 3,064 candidate genes have glutamine at this position, all 40 genes whose large red-shift gains were suggested by our ML-based model have amino acids other than glutamine, which suggests that our ML-based model avoided the genes having glutamine at position 93. Especially, 12 (37.5%) of 32 genes that actually showed red-shifted absorption compared with the base wavelengths had methionine at this position (Extended Data Figure 2), which is substantially higher than the proportion of methionine-conserving genes in the 3,064 candidates (16.1%). The red-shifting effect of the L-to-M mutation of this residue in GPRs previously reported33 and the current result imply that many rhodopsins have evolved methionine to absorb light with longer wavelengths. Position 215 in BR is also known to have a colour-tuning role. The mutation from alanine to threonine or serine (A/TS switch) has a blue-shifting effect of 9–20 nm^16,34–36^. Five of seven genes that showed blue-shifted *λ*_max_ compared with the base wavelengths have threonine or serine at this position, suggesting that these types of genes should be avoided to explore red-shifted rhodopsins. By contrast, asparagine was conserved in more than half (58.4%) of the 3,064 candidate genes, especially in those belonging to the PR subfamily. A significant portion (37.5%) of the genes with red-shifted absorption compared with the base wavelengths also had asparagine at this position (Extended Data Figure 2). The A-to-N mutation at this position had a smaller effect (4–7 nm)^20,35^ than that of the A-to-S/T mutation; thus, the difference between alanine and asparagine is not so critical to explore red-shifted rhodopsins. Position 186 in BR is proline in most microbial rhodopsins (in 98.7% of the 3,064 candidate genes), and the mutation to non-proline amino acids induces red-shift of absorption^16^. We identified sodium pump rhodopsin (NaR) from *Parvularcula oceani*, which also has a threonine at this position, and showed 10-nm longer absorption than the base wavelength. Although genes having non-proline amino acids are rare in nature, it would be beneficial to identify new red-shifted rhodopsins. These results indicate that ML-based modelling can provide insights for identifying new functional tuning rules for proteins based on specific amino acid residues.

The number of reported microbial rhodopsin genes is rapidly increasing because of the development of next-generation sequencing techniques and microbe culturing methods. New microbial rhodopsins with molecular characteristics suitable for optogenetics applications are expected to be included in upcoming genomic data. Our ML-based model could be expected to reduce the costs associated with identifying red-shifted rhodopsins from these data.

Especially, we expect that our ML-based model could be applied to ion channel and enzymatic rhodopsins, which were not a focus of this study because of their eukaryotic origins; however, their use in optogenetics research could help identify more useful optogenetics tools with red-shifted absorption in the future.

## Methods

### Construction of training and target data sets

In this study, we constructed a new training data set (Extended Data Table 1) by adding 97 genes for which the *λ*_max_ had recently been reported in the literature or determined by our experiments, to a previously reported data set^20^. The sequences were aligned using ClustalW^37^ and the results were manually checked to avoid improper gaps and/or shifts in the TM parts. The aligned sequences were then used for ML-based modeling.

To collect microbial rhodopsin genes for the training data set, BR^38^ and heliorhodopsin 48C12^39^ sequences were used as queries for searching homologous amino acid sequences in NCBI non-redundant protein sequences and metagenomic proteins^27^ and the *Tara* Oceans microbiome and virome database^28^. Protein BLAST (blastp)^26^ was used for the homology search, with the threshold E-value set at < 10 by default, and sequences with > 180 amino acid residues were collected. All sequences were aligned using ClustalW37. The highly diversified C-terminal 15-residue region behind the retinal binding Lys (BR K216) and long loop of HeR between helices A and B were removed from the sequences to avoid unnecessary gaps in the alignment. The successful alignment of the TM helical regions, especially the 3rd and 7th helices, was checked manually. The phylogenic tree was drawn using the neighbor-joining method^40^, and the microbial rhodopsin subfamilies were categorized based on the phylogenetic distances, as reported previously^29^. Based on the phylogenetic tree, 3,064 putative ion-pumping rhodopsin genes from bacterial and archaeal origins were extracted, and their aligned sequences were used as the training data set for the prediction of *λ*_max_.

### ML modeling

Suppose that we have *K* pairs of an amino acid sequence and an absorption wavelength 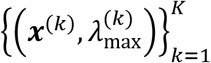, where ***x***^(k)^ ∈ ℝ^*MN*^ is the feature vector of the fc-th amino-acid sequence and 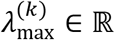 is the absorption wavelength of the *K*-th rhodopsin protein. The least-absolute shrinkage selection operator (LASSO) is a standard regression model in which important regression coefficients can be automatically selected by the penalty on the absolute value of the coefficient, as follows:

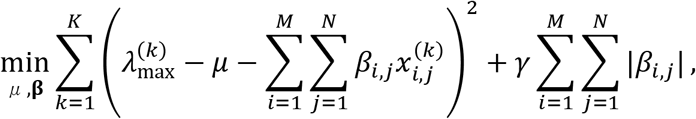

where ***β*** ∈ ℝ^*MN*^ is a vector of *β_i,j_* and *γ* > 0 is the regularization parameter. BLASSO is a Bayesian extension of LASSO for which the model is defined through the following random variables:

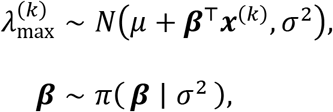

where *N*(*μ*, *s*^2^) is a Gaussian distribution with mean *μ*; and variance *s*^2^, and 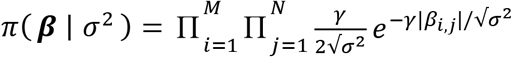 is the conditional Laplace prior. In this model, the maximum of the conditional distribution of the parameter 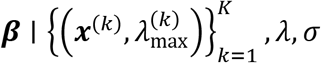 is equivalent to the LASSO^41^ estimator. For the computational details, see the original paper^25^. Since the resulting predictive distribution of *f*(***x***) is not analytically tractable, the parameters *β* and *μ* are sampled from the estimated distribution *T* = 10,000 times. For each candidate ***x***, we approximately obtain 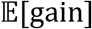 by

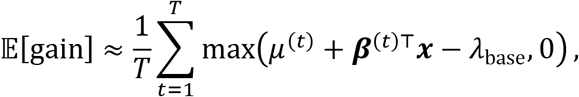

where *μ*^(*t*)^ and *β*^(*t*)^ are the *t*-th sampled parameters.

### Protein expression

The synthesized genes of microbial rhodopsins codon-optimized for *E. coli* (Genscript, NJ) were incorporated into the multi-cloning site in the pET21a(+) vector (Novagen, Merck KGaA, Germany). The plasmids carrying the microbial rhodopsin genes were transformed into the *E. coli* C43(DE3) strain (Lucigen, WI). Protein expression was induced by 1 mM isopropyl β-D-1-thiogalactopyranoside (IPTG) in the presence of 10 μM *all-trans* retinal for 4 h.

### Measurement of the absorption spectra and *λ*_max_ of rhodopsins by bleaching with hydroxylamine

*E. coli* cells expressing rhodopsins were washed three times with a solution containing 100 mM NaCl and 50 mM Na2HPO4 (pH 7). The washed cells were treated with 1 mM lysozyme for 1 h and then disrupted by sonication for 5 min (VP-300N; TAITEC, Japan). To solubilize the rhodopsins, 3% *n*-dodecyl-D-maltoside (DDM, Anatrace, OH) was added and the samples were stirred for overnight at 4 °C. The rhodopsins were bleached with 500 mM hydroxylamine and subjected to yellow light illumination (λ > 500 nm) from the output of a 1-kW tungsten–halogen projector lamp (Master HILUX-HR; Rikagaku) through coloured glass (Y-52; AGC Techno Glass, Japan) and heat-absorbing filters (HAF-50S-15H; SIGMA KOKI, Japan). The absorption change upon bleaching was measured by a UV-visible spectrometer (V-730; JASCO, Japan).

### Ion-transport assay of rhodopsins in *E. coli* cells

To assay the ion-transport activity in *E. coli* cells, the cells carrying expressed rhodopsin were washed three times and resuspended in unbuffered 100 mM NaCl. A cell suspension of 7.5 mL at OD660 = 2 was placed in the dark in a glass cell at 20 °C and illuminated at *λ* > 500 nm from the output of a 1 kW tungsten-halogen projector lamp (Rikagaku, Japan) through a long-pass filter (Y-52; AGC Techno Glass, Japan) and a heat-absorbing filter (HAF-50S-50H; SIGMA KOKI, Japan). The light-induced pH changes were measured using a pH electrode (9618S-10D; HORIBA, Japan). All measurements were repeated under the same conditions after the addition of 10 μM CCCP.

### Reporting Summary

Further information on experimental design is available in the Nature Research Reporting Summary linked to this article.

## Data Availability

Data supporting the findings of this manuscript are available from the corresponding author upon reasonable request.

## Acknowledgments

This work was supported by Grants-in-Aid from the Japan Society for the Promotion of Science (JSPS) for Scientific Research (KAKENHI grant Nos. 17H03007 to K.I., 17H04694 and 16H06538 to M.Karasuyama, 19H04959 to H.K., and 17H00758 and 16H06538 to I.T.), the Japan Science and Technology Agency (JST), PRESTO, Japan (grant Nos. JPMJPR15P2 to K.I. and JPMJPR15N2 to M.Karasuyama), and CREST, Japan (grant No. JPMJCR1502) to I.T.; K.I., H.K., and I.T. received support from RIKEN AIP; O.B. received support from the Louis and Lyra Richmond Memorial Chair in Life Sciences.

## Author contributions

K.I., R.G., O.B., and H.K. contributed to the study design; K.I., D.Y., K.Y., and O.B. collected sequences of non-redundant and metagenomic rhodopsin genes from the GenBank and *Tara* Oceans metagenomic data sets and conducted multiple amino-acid alignments of rhodopsins; M.Karasuyama, Y.I., and I.T. constructed the machine learning method to estimate the absorption wavelengths of microbial rhodopsins; R.N. constructed DNA plasmids of microbial rhodopsins and introduced them into *E. coli* cells; R.N., K.M., and T.N. conducted expressions of microbial rhodopsins in *E. coli* cells and determined their *λ*_max_ by hydroxylamine bleaching; M.Konno carried out the ion-transport assay of rhodopsins in *E. coli* cells; K.I., M.Karasuyama., H.K., and I.T. wrote the paper; All authors discussed and commented on the manuscript.

## Supporting Information

**Extended Data Figure 1.**
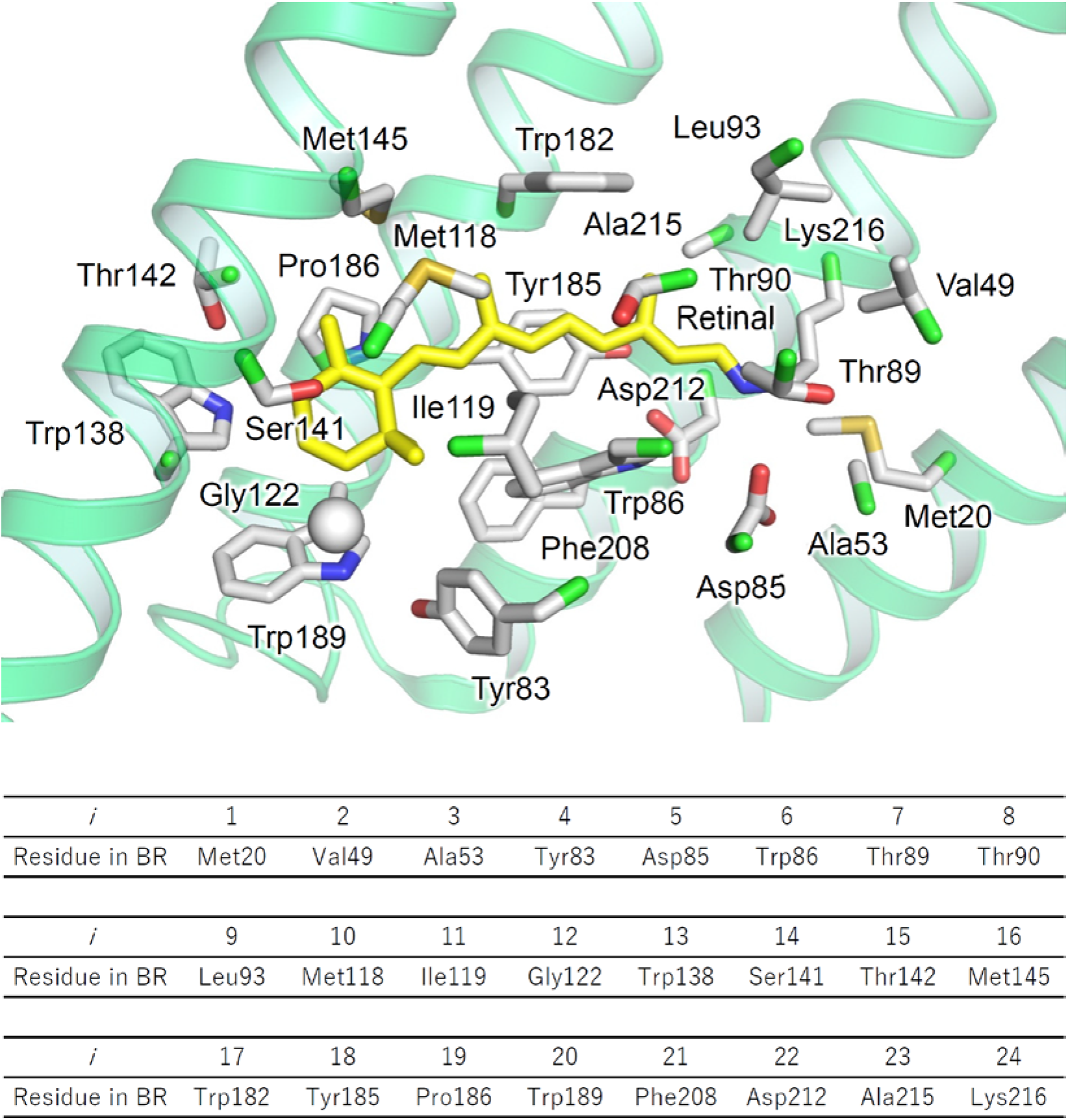
Amino acid residues around the retinal chromophore. The structure of the 24 amino acid residues around the retinal used in the current ML model in the X-ray crystallographic structure of BR (PDB ID: 1IW6 (Matsui et al. *J. Mol. Biol*. (2002) **324**, pp. 469–481)). The Cα atom of Gly122 is shown as a white sphere. For clarity, the ribbon models of helices B, C, and E were omitted. The table lists the residue numbers and names of each residue in BR.

**Extended Data Figure 2.**
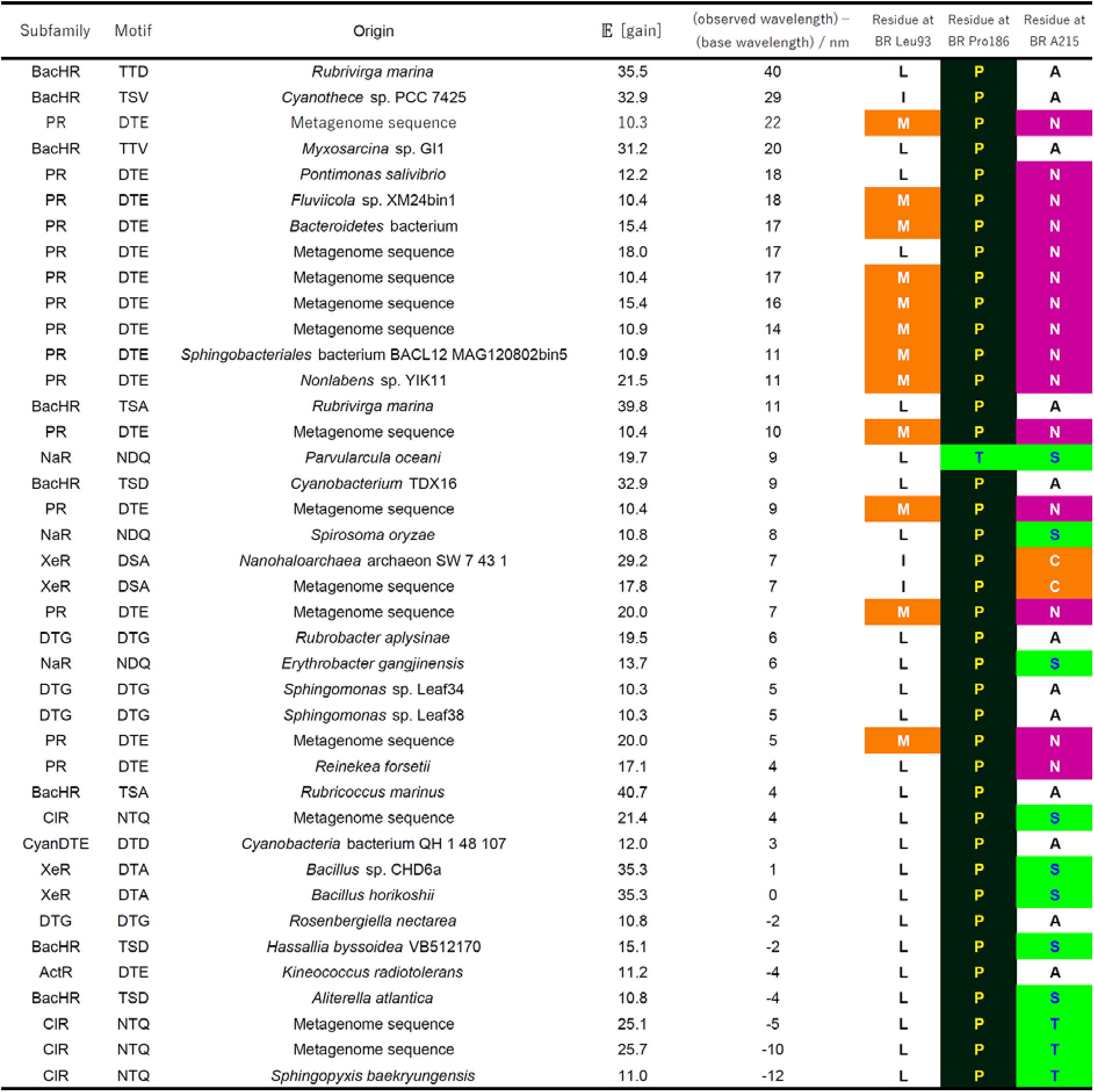
Amino acid residues at the color-tuning positions. The amino acid residues at the color-tuning positions corresponding to BR Leu93, Pro189, and Ala215.

## References

1 Ernst, O. P. et al. Microbial and animal rhodopsins: Structures, functions, and molecular mechanisms. Chem. Rev. 114, 126–163 (2014).

2 Oesterhelt, D. & Stoeckenius, W. Rhodopsin-like protein from the purple membrane of *Halobacterium halobium*. Nat. New Biol. 233, 149–152 (1971).

3 Oesterhelt, D. & Stoeckenius, W. Functions of a new photoreceptor membrane. Proc. Natl. Acad. Sci. USA 70, 2853–2857 (1973).

4 Man, D. et al. Diversification and spectral tuning in marine proteorhodopsins. EMBO J. 22, 1725–1731 (2003).

5 Venter, J. C. et al. Environmental genome shotgun sequencing of the sargasso sea. Science 304, 66–74 (2004).

6 Inoue, K., Kato, Y. & Kandori, H. Light-driven ion-translocating rhodopsins in marine bacteria. Trends. Microbiol. 23, 91–98 (2014).

7 Inoue, K. et al. A light-driven sodium ion pump in marine bacteria. Nat. Commun. 4, 1678 (2013) 10.1038/ncomms2689.

8 Nagel, G. et al. Channelrhodopsin-1: A light-gated proton channel in green algae. Science 296, 2395–2398 (2002).

9 Niho, A. et al. Demonstration of a light-driven SO_4_^2−^ transporter and its spectroscopic characteristics. J. Am. Chem. Soc. (2017).

10 Deisseroth, K. Optogenetics: 10 years of microbial opsins in neuroscience. Nat. Neurosci. 18, 1213–1225 (2015).

11 Liu, X. et al. Optogenetic stimulation of a hippocampal engram activates fear memory recall. Nature 484, 381–385 (2012).

12 Ramirez, S. et al. Creating a false memory in the hippocampus. Science 341, 387–391 (2013).

13 Yizhar, O. et al. Neocortical excitation/inhibition balance in information processing and social dysfunction. Nature 477, 171–178 (2011).

14 Marshel, J. H. et al. Cortical layer-specific critical dynamics triggering perception. Science 365, eaaw5202 (2019).

15 Schneider, F., Grimm, C. & Hegemann, P. Biophysics of channelrhodopsin. Annu. Rev. Biophys. 44, 167–186 (2015).

16 Inoue, K. et al. Red-shifting mutation of light-driven sodium-pump rhodopsin. Nat. Commun. 10, 1993 (2019).

17 Ganapathy, S. et al. Retinal-based proton pumping in the near infrared. J. Am. Chem. Soc. 139, 2338–2344 (2017).

18 Pushkarev, A. et al. The use of a chimeric rhodopsin vector for the detection of new proteorhodopsins based on color. Front. Microbiol. 9, 439 (2018).

19 Oda, K. et al. Crystal structure of the red light-activated channelrhodopsin Chrimson. Nat. Commun. 9, 3949 (2018) 10.1038/s41467-018-06421-9.

20 Karasuyama, M., Inoue, K., Nakamura, R., Kandori, H. & Takeuchi, I. Understanding colour tuning rules and predicting absorption wavelengths of microbial rhodopsins by data-driven machine-learning approach. Sci. Rep. 8, 15580 (2018).

21 Pedraza-González, L., De Vico, L., Marín, M. d. C., Fanelli, F. & Olivucci, M. A-arm: Automatic rhodopsin modeling with chromophore cavity generation, ionization state selection, and external counterion placement. J. Chem. Theory Comput. 15, 3134–3152 (2019).

22 Bishop, C. M. Pattern recognition and machine learning. (Springer, 2006).

23 Snoek, J., Larochelle, H. & Adams, R. P. in Advances in Neural Information Processing Systems 25 (NIPS 2012). (eds F. Pereira, C. J. C. Burges, L. Bottou, & K. Q. Weinberger) 2951–2959 (Curran Associates, Inc.).

24 Shahriari, B., Swersky, K., Wang, Z., Adams, R. P. & Freitas, N. d. in Proceedings of the IEEE. 148–175.

25 Park, T. & Casella, G. The bayesian lasso. J. Am. Stat. Assoc. 103, 681–686 (2008).

26 Johnson, M. et al. Ncbi blast: A better web interface. Nucleic Acids Res. 36, W5–W9 (2008).

27 Brown, G. R. et al. Gene: A gene-centered information resource at ncbi. Nucleic Acids Res. 43, D36–D42 (2015).

28 Sunagawa, S. et al. Ocean plankton. Structure and function of the global ocean microbiome. Science 348, 1261359 (2015).

29 Yamauchi, Y. et al. Engineered functional recovery of microbial rhodopsin without retinal-binding lysine. Photochem Photobiol 95, 1116–1121 (2019).

30 Hasemi, T., Kikukawa, T., Kamo, N. & Demura, M. Characterization of a cyanobacterial chloride-pumping rhodopsin and its conversion into a proton pump. J. Biol. Chem. 291, 355–362 (2016).

31 Harris, A. et al. Molecular details of the unique mechanism of chloride transport by a cyanobacterial rhodopsin. Phys. Chem. Chem. Phys. 20, 3184–3199 (2018).

32 Béjà, O. et al. Bacterial rhodopsin: Evidence for a new type of phototrophy in the sea. Science 289, 1902–1906 (2000).

33 Ozaki, Y., Kawashima, T., Abe-Yoshizumi, R. & Kandori, H. A color-determining amino acid residue of proteorhodopsin. Biochemistry 53, 6032–6040 (2014).

34 Shimono, K., Ikeura, Y., Sudo, Y., Iwamoto, M. & Kamo, N. Environment around the chromophore in *pharaonis* phoborhodopsin: Mutation analysis of the retinal binding site. Biochim. Biophys. Acta 1515, 92–100 (2001).

35 Sudo, Y. et al. A blue-shifted light-driven proton pump for neural silencing. J. Biol. Chem. 288, 20624–20632 (2013).

36 Inoue, K. et al. Converting a light-driven proton pump into a light-gated proton channel. J. Am. Chem. Soc. 137, 3291–3299 (2015).

37 Thompson, J. D., Higgins, D. G. & Gibson, T. J. Clustal-W - improving the sensitivity of progressive multiple sequence alignment through sequence weighting, position-specific gap penalties and weight matrix choice. Nucleic Acids Res. 22, 4673–4680 (1994).

38 Khorana, H. G. et al. Amino acid sequence of bacteriorhodopsin. Proc. Natl. Acad. Sci. USA 76, 5046–5050 (1979).

39 Pushkarev, A. et al. A distinct abundant group of microbial rhodopsins discovered using functional metagenomics. Nature 558, 595–599 (2018).

40 Saitou, N. & Nei, M. The neighbor-joining method: A new method for reconstructing phylogenetic trees. Mol. Biol. Evol. 4, 406–425 (1987).

41 Tibshirani, R. Regression shrinkage and selection via the lasso. Journal of the Royal Statistical Society. Series B (Methodological) 58, 267–288 (1996).

